# Flux estimation analysis systematically characterizes the metabolic shifts of the central metabolism pathway in human cancer

**DOI:** 10.1101/2022.10.27.514080

**Authors:** Grace Yang, Shaoyang Huang, Kevin Hu, Alex Lu, Jonathan Yang, Noah Meroueh, Pengtao Dang, Haiqi Zhu, Sha Cao, Chi Zhang

**Author notes:** Equal contribution (Grace Yang, Shaoyang Huang, Kevin Hu, Alex Lu).

## Abstract

Glucose and glutamine are major carbon and energy sources that promote the fast proliferation of cancer cells. Metabolic shifs observed on cell line or mouse models may not reflect the general metabolic shifts in real human cancer tissue. In this study, we conducted a computational characterization of the flux distribution and variations of the central energy metabolism and key branches in a pan-cancer analysis, including glycolytic pathway, production of lactate, TCA cycle, nucleic acids synthesis, glutaminolysis, glutaminate, glutamine and glutathione metabolism, amino acids synthesis, in 11 cancer subtypes and 9 matched adjacent normal tissue types, by using TCGA tissue transcriptomics data. Our analysis confirms the increased influx in glucose uptake and glycolysis and decreased upper part of TCA cycle, i.e., Warbug effect in the analyzed cancer types. However, consistently increased lactate production and second half of TCA cycle were only seen in certain cancer types. More interestingly, we did not see cancer tissues have highly shifted glutaminolysis compared to their adjacent normal controls. A systems biology model of metabolic shifts in through cancer and tissue types is further developed and analyzed. We observed that (1) normal tissues have distinct metabolic phenotypes, (2) cancer types have drastically different metabolic shifts compared to their adjacent normal controls, and (3) the different shifts happened to tissue specific metabolic phenotypes result in a converged metabolic phenotype through cancer types and cancer progression. This study strongly suggests the possibility to have a unified framework for studies of cancer-inducing stressors, adaptive metabolic reprogramming, and cancerous behaviors.

## INTRODUCTION

Dysregulation of metabolic pathways is a hallmark cancer. In the past decades, new biotechnologies and experimental systems advanced substantial knowledge of metabolic shifts and their functional roles of in the oncogenesis process and the progression of cancer. Metabolic phenotypes of cancer and stromal cells and mechanistic insights of how the metabolic system is shifted along the co-evolution of cancer and the tumor microenvironment (TME) have been discovered on different experimental systems, such as cancer cell line, mouse model, patient derived xenografts or organoid models. Despite a plethora of knowledge on the core components of metabolic pathways we have gained, there are still major gaps in our understanding of the integrated behavior and metabolic heterogeneity of cells in cancer microenvironment. Essentially, the metabolic behavior can be determined by different factors and vary dramatically from cell to cell and tissue to tissue due to their high plasticity, driven by the need to cope with various dynamic metabolic requirements and biochemical conditions.

Warburg effect, characterized by high glycolytic activity and decreased TCA cycle, is considered as a common metabolic reprogramming mechanism in human solid cancer [1]. The discovery of glutaminolysis elucidate the role of “fueling” for energy production and biosynthesis of other metabolites from amino acids, which also expanded the definition of central metabolism in cancer metabolism. In addition to these “common” metabolic shifts, variations in branches of the central metabolic pathway have been observed in different cancer types, including biosynthesis of nucleotide, biosynthesis of serine and glycine, Cori cycle and gluconeogenesis, malate shuttle and aspartate metabolism, redox balance and glutathione metabolism, biosynthesis of fatty acids, synthesis of immune-metabolite 2-hydroxyglutarate from a-ketoglutarate, cytosolic metabolism of glutamine and glutamate. Noted, majority of these analysis are made on cell line or mouse systems, which cannot mimic the dysbalanced redox, pH and oxygen levels. In addition, nutrient supplies of the experimental system also differ from a real cancer tissue, such as both glucose and glutamine are always sufficiently provided under experimental conditional while their availability level and ratios heavily shifts through cancer and determines metabolic phenotypes. In addition, recent spatial techniques suggest the heterogenous distribution of metabolic stresses in a real cancer tissue, which promote metabolic competition and co-adaptation between cancer and stromal cells. All the evidence suggests that the experimental systems under normal physiological conditions have drastically different biochemical properties compared to a real human cancer TME.

We would like to point out the main observations from experimental systems may unnecessarily reflect the metabolic and nutrient-partition activities in human cancer tissues. For example, a recent study presented that myeloid cells consume the highest amount of glucose per cell in mouse tumor tissues, followed by T cells and tumor cells [2]. A simple analysis by correlating predicted immune cell population with metabolic gene expression levels could eliminate this possibility in real TME (see supplementary analysis). Indeed, based on transcriptomic data analyses of human cancer and transplant tissues as well subcutaneous mouse tumors, that the main observations reported in the article do not reflect the immune and nutrient-partition activities in human cancer tissues, instead they resemble the relevant activities in human transplants. Although numerous analyses of metabolic variations have been conducted on omics data collected from cancer tissue samples to the best of our knowledge, there is a lack of explicit analysis that mechanically estimate and quantify metabolic shifts. We have recently developed a new graph neural network-based method, namely single cell flux estimation analysis (scFEA), to predict metabolic flux by using single cell transcriptomics data. scFEA computes sample-wise

To provide a less biased and comprehensive characterization of the landscape of metabolic changes in in human solid cancer, we conduct a systematic evaluation of the metabolic reprogramming and characteristics via a computational analysis on pan-cancer transcriptomics data. Specifically, we first reconstructed the central metabolism pathway by including glucose, glutamine, and glutathione metabolism and six branches of the central metabolism network in a subcellular resolution. We modified our inhouse developed metabolic fluxome predictor, namely single-cell Flux Estimation Analysis (scFEA) to enable the analysis on TCGA pan-cancer tissue transcriptomics data. Our analysis revealed distinct metabolic variations and shifts through different cancer types. As Warburg effect has been identified in almost all analyzed cancer tissues, we did not see a significant contribution of glutaminolysis in fueling the TCA cycle. We identified that (1) normal tissues have distinct metabolic phenotypes, (2) cancer types have drastically different metabolic shifts, and (3) the different shifts happened to tissue specific metabolic phenotypes result in a converged metabolic phenotype through cancer types and cancer progression. Our analysis brought novel insights into the understanding of metabolic shifts in human cancer. The cancer and tissue type specific metabolic shifts and the resulted convergent metabolic phenotype suggested the necessity of personalized therapeutic strategy or nutrient and diet design for targeting metabolism in cancer treatment.

## RESULTS

### Reconstruction of central metabolism pathway in the sub-cellular resolution

To comprehensively evaluate the variations of energy metabolism in cancer, we collect the central metabolism and its branches from KEGG database and manually curate the reaction information from literature data. As subcellular compartments have different levels of enzymes, substrates, biochemical characteristics, and kinetic parameters, sub-cellular localization information of reactions is needed to accurately assess their stoichiometric relations. **Figure 1B** illustrates the reconstructed central metabolism network, including 42 reaction modules, 27 intermediate metabolites, 15 end metabolites, and 320 genes in cytosol, mitochondria, and extracellular regions. The reconstructed central metabolism network include six major pathways, namely glycolysis, upper and lower parts of TCA cycle, glutaminolysis, glutamine and glutamate metabolism, and glutathione metabolism, and six other branches, namely Glyceraldehyde 3-phosphate (G3P) to nucleotide synthesis, 3-Phospho-D-glycerate (3PD) to serine synthesis, aspartate-malate shuttle, mitochondrial citrate fueling of fatty acid synthesis, transport of 2-Oxoglutarate (2OG) to cytosol, and transform of 2OG to 2-Hydroxyglutarate (2HG).

**Figure 1.**
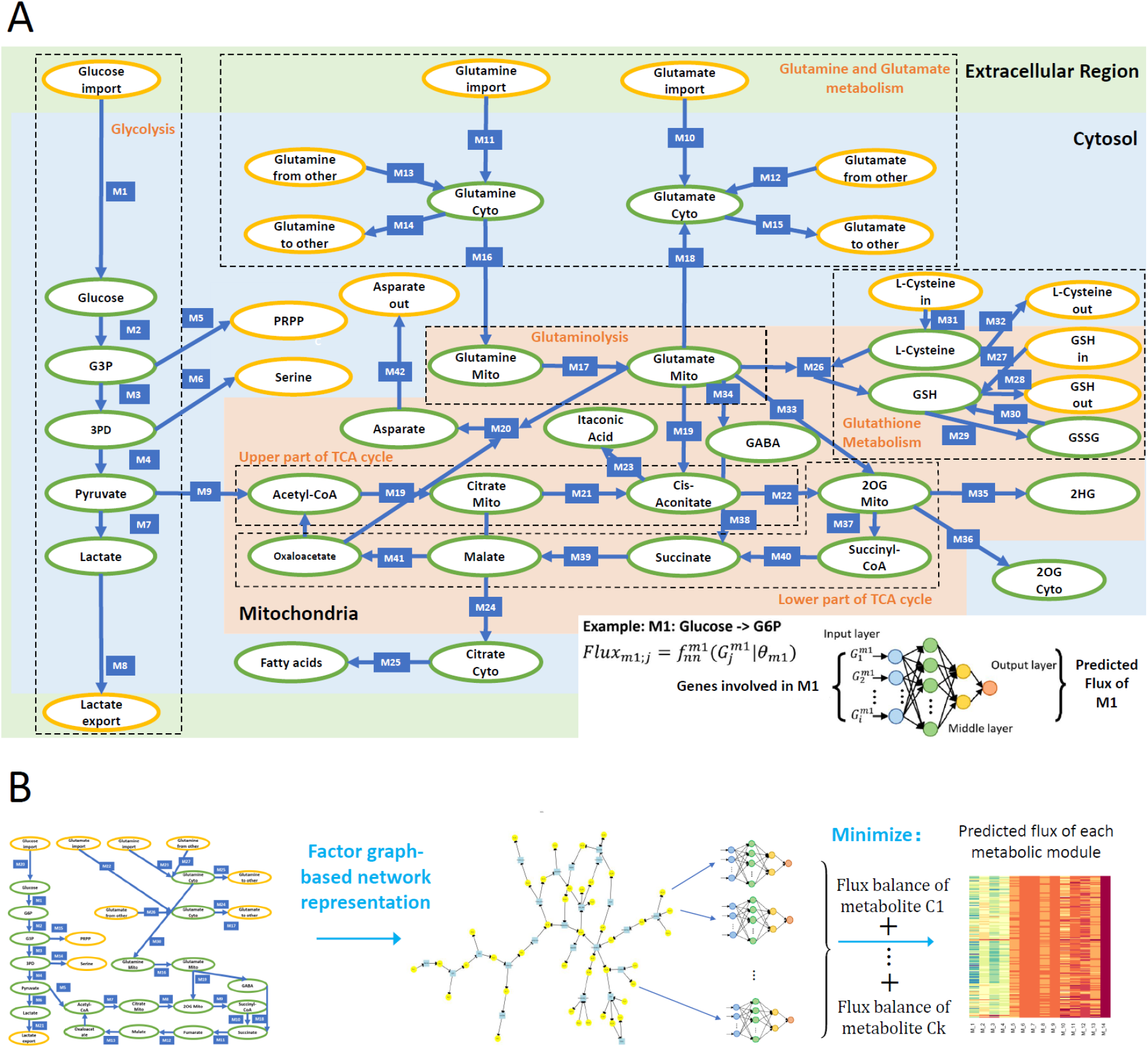
**(A) Reconstructed Central Metabolism Pathwa**y. Intermediate and end metabolites are green and yellow labeled. The example on the bottom right showcases the neural network-based flux predictive model, which takes genes involved in module 1 (M1) and output predicted flux. **(B) Analysis framework of scFEA.**

### Estimation of sample-wise metabolic flux and metabolite abundance of central metabolism

We modified our recently developed method, scFEA, to enable the application on tissue transcriptomics data [2] (see details in methods). scFEA models metabolic fluxes in a tissue based on gene expression data of a large number of samples by assuming (1) the total influx of each metabolite is approximately the same as its total outflux, which is constrained by a quadratic loss over the metabolic network; and (2) changes in the rate of each reaction can be modeled as a (non-linear) function of changes in the expression levels of genes involved in the reaction and its neighbors, which could be modeled by a neural network. Note that assumption (1) is generally true unless some major in-/out-flux for a metabolite is not considered. Assumption (2) is a combination of three simpler assumptions: (1) the concentration of an enzyme is a function of reaction rate that could be well supported by Michaelis Menton equation; (2) the concentration of an enzyme is also a (non-linear) function of the expression level of its encoding genes, with both functions being invariant across different samples of different cancer types; and (3) there exists a latent non-linear relation between the concentration of metabolites and the genes involved in its transport and relevant reactions. Both assumptions (1) and (2) are supported by published studies [3–5]. Intuitively, one can think this model (for each reaction) as an integrated Michaelis–Menten model, whose parameters and non-linear form are implicitly estimated by a neural network using the large number of available gene expression data (**Figure 1A**). Detailed formulation and parameters of the scFEA method utilized in this study is available in Methods.

**Figure 1B** outlines the flowchart of scFEA with further details given in Materials and Methods. scFEA models the metabolic flux of each module by a three layer fully connected neural network of genes involved in the module, which minimizes the total imbalance of the intermediate substrates across all tissue samples. For the central metabolism network with *K* = 42 modules and # genes as the average number of genes encoded in each reaction module, there are 12 × *K* × (# genes) unknowns to be estimated, where 12 is determined by the neural network architecture. On the other side, there are *K×N* constraints, where *K* = 27 and *N* = 5253 are the numbers of intermediate substrates and samples, respectively. As *K*×*N*» 12 × *K* × (# genes) in this study, the large number of samples in the utilized TCGA pan-cancer data enables sufficient statistical power to reliably estimation of the unknowns.

Here we first validated scFEA on a simplified central metabolic network by applying the method on our recently collected scRNA-seq data of 168 patient-derived pancreatic cancer cell lines Pa03c under four conditions: normoxia (N), hypoxia (H), normoxia and knock-down of APEX1 (N-APEX1-KD), and hypoxia and knock-down of APEX1 (H-APEX1-KD) [2]. APEX1 plays a central role in the cellular response to oxidative stress [6]. As scFEA has been previously validated over a simplified central metabolism data, the goal of this validation is the confirm that scFEA can capture the metabolic changes on the newly reconstructed central metabolism pathway. scFEA predicts increased level of TCA cycle and decreased glycolysis in normoxic cells than hypoxic cells and increased glutathione (GSH) to Glutathione disulfide (GSSG) in APEX knockout cells. These observations match our recent studies experimentally confirmed metabolic changes in the Pa03c cells, including (1) the knockdown of APEX1 results in increased oxidative stress and cell death, (2) hypoxia triggered increased glycolytic activity and lactate production, (3) decreased TCA cycle, and (4) insignificantly changed glutamine metabolism in Pa03c cells [7]. These observations demonstrated that the scFEA prediction can capture the major variations in the reconstructed central metabolism network under different biochemical conditions.

We further applied the modified scFEA on a TCGA pan-cancer transcriptomics data of 11 cancer and sub-cancer types having matched adjacent normal controls, namely breast cancer (BRCA) luminal, her2- and triple negative (TNBC) subtypes, colon adenocarcinoma (COAD), head and neck cancer (HNSC), kidney renal clear cell carcinoma (KIRC), kidney renal papillary cell carcinoma (KIRP), lung adenocarcinoma (LUAD), prostate adenocarcinoma (PRAD), stomach adenocarcinoma (STAD), and thyroid carcinoma (THCA), totaling 5253 samples (see details in methods). Noted, scFEA predicts sample wise in-/out-flux for each intermediate metabolite. Hence, the relative abundance change of the intermediate metabolites can be estimated by the difference of their in- and out-flux in each sample.

### A pan-cancer level evaluation of metabolic variations of the central metabolic pathway and its branches

We first evaluate the flux changes between cancer and adjacent normal tissue. Our analysis suggested that the flux of glycolytic pathway, lactate production, lactate export, biosynthesis of glutathione, glutamate and glutamine import, and glutamine, glutamate, and aspartate metabolism to other amino acids in cytosol are consistently increased in cancer vs normal in all or almost all examined cancer types (**Figure 2A-D**, **Supplementary Figure S1**). We also observed consistently decreased flux of fatty acid biosynthesis, glutathione to other amino acids, enzyme catalyzed glutathione to GSSG, and cysteine metabolism (**Figure 2E-F, Supplementary Figure S1**). The module from citrate to 2OG is consistently decreased while the oxaloacetate to citrate is increased in the upper part of TCA cycle. The variations of the low part of TCA cycle are differed through cancers. Although the glutamine and glutamate metabolism show distinct variations in cancer vs normal, we did not see a significant difference in glutaminolysis, i.e., glutamate to 2OG or glutamate to GABA and succinate in mitochondria. We also did not observe a significant increase in the low part of TCA cycle. Most cancer types tend to have decrease fumarate to malate and malate to oxaloacetate. **Supplementary Figure S1** illustrates detailed flux changes of metabolic modules in the reconstructed central metabolism network.

**Figure 2.**
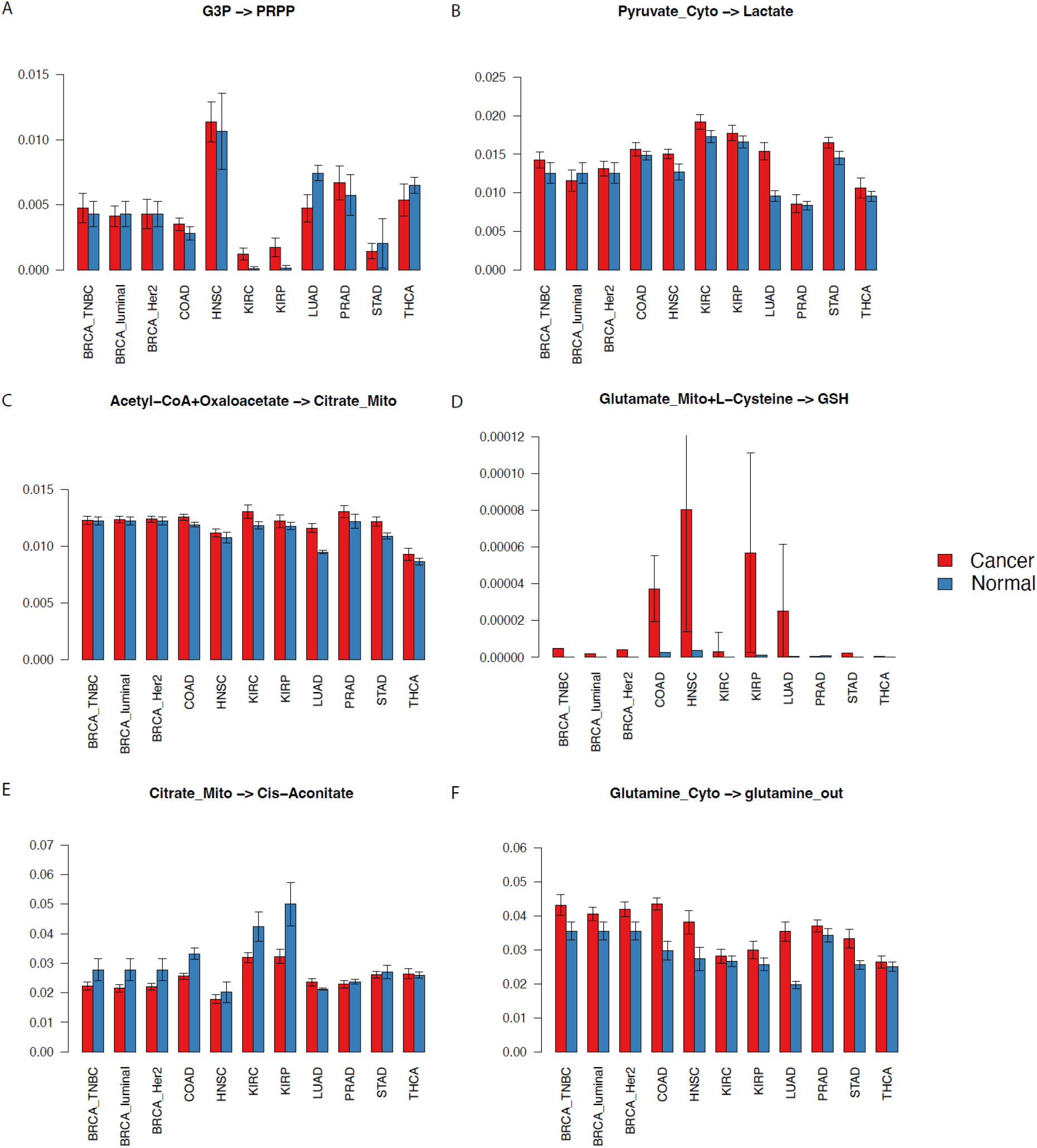
Selected fluxomic changes in cancer vs normal. X-axis and y-axis represent cancer types and predicted flux, respectively.

We also checked the predicted metabolomic changes in the central metabolism network. Noted, total flux balance assumption was made in scFEA by a quadratic loss. The metabolomic change reflect a trend of the variation of metabolites rather than their exact concentration. We predicted consistently depleted glycolytic and TCA cycle intermediates, mitochondrial glutamine, and cytosolic glutamate in cancer vs normal while the abundance of lactate, aspartate, succinyl-CoA, and cytosolic glutamine tend to be increased. **Figure 3** shows the detailed metabolic variations identified in this study. Below we discuss potential mechanistic interpretation of the observed metabolic variations.

**Figure 3.**
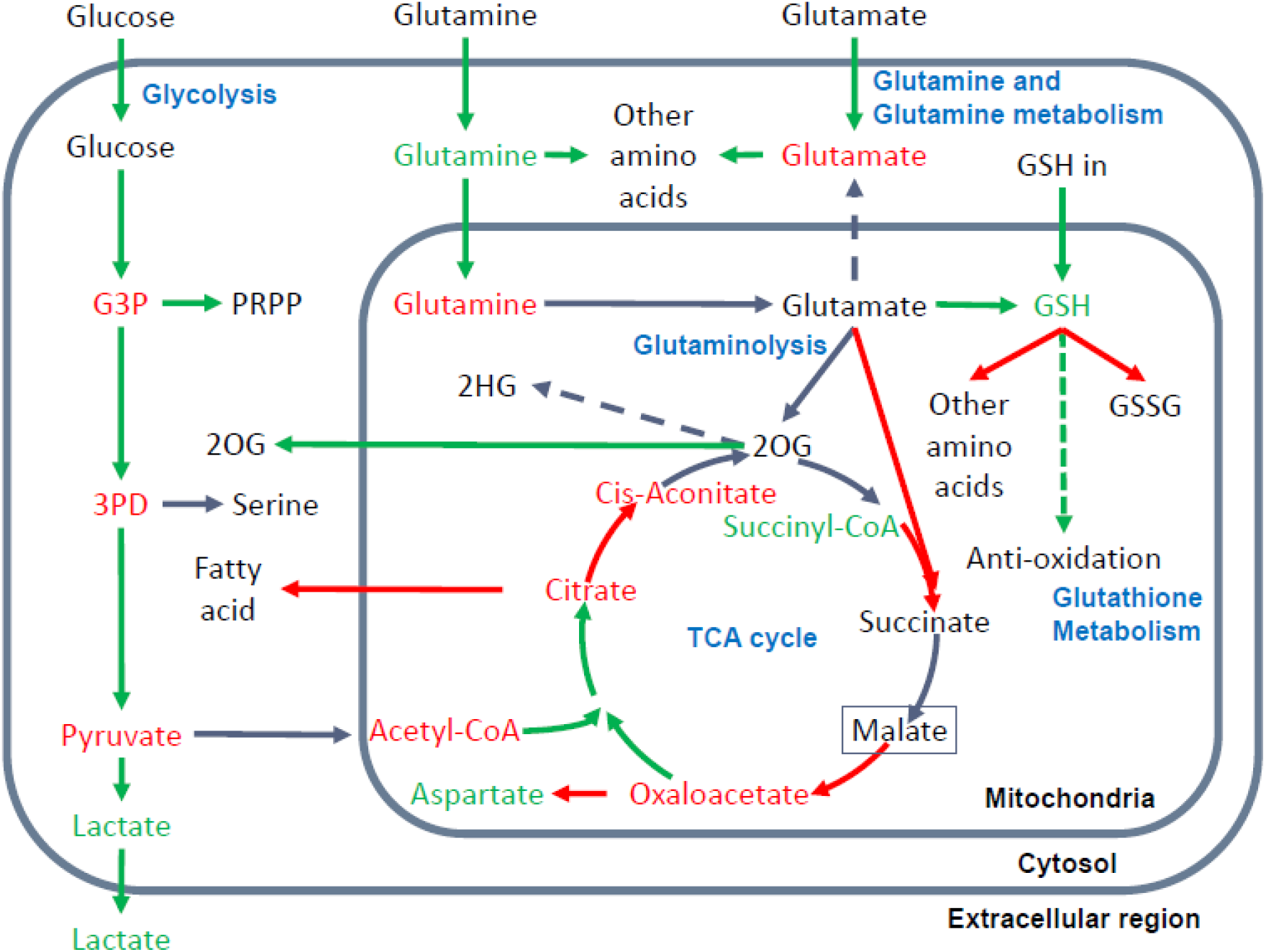
Metabolic variations in central metabolism network observed in this study. Increased or decreased flux of a reaction module is represented by a green or red arrow; accumulation or depletion of metabolites are green and red colored; the reactions and metabolites of inconsistent changes are grey and black colored, respectively. Dash arrow suggests the predict flux of the reaction is consistently zero or substantially low. The green dash arrow of GSH to anti-oxidation suggests our inferred increased flux of GSH in anti-oxidation in cancer.

Glycolysis: We observed increased glycolytic flux, depleted intermediates, and increased lactate production in almost all examined cancer types. These observations are consistent with well discussed Warburg effect. The only contradiction to experimentally reported result is that we did not see a significant increase in glucose uptake. Noted, the flux prediction is based on both gene expression and neighboring fluxes. Our previous studies confirmed the consistently up regulated glucose transporter. The scFEA computes the real glucose uptake is not increased, probably because the availability of glucose in TME is limited.

TCA cycle: We observed that majority of the TCA cycle is decreased in cancer vs normal, except for the first step of acetyl-coA to citrate. The rate limiting step from citrate to cis-aconitate is consistently decreased and most cancer types have decreased succinate to malate and malate to oxaloacetate. We also observed consistent decrease of TCA cycle intermediates, except for the increased succinyl-CoA and varied changes of 2OG, succinate and malate through cancer types. Our explanation is that the overall TCA cycle is suppressed but the fueling from glutaminolysis relieved the depletion of TCA cycle intermediates at a certain level. However, in the TME of human cancer, the flux of carbon source from glutaminolysis is not enough to revert the decreased low part of TCA cycle.

Glutamate and glutamine metabolism: The uptake of glutamate and glutamine is consistently increased in cancer vs normal. However, our flux analysis suggests that majority of the glutamate and glutamine are utilized for biosynthesis of other amino acids in cytosol rather than transport into mitochondria to fuel TCA cycle. We saw the transport of glutamate from mitochondria to cytosol is almost zero in all cancer types. Majority of the mitochondrial glutamate is utilized for glutathione biosynthesis than glutaminolysis.

Glutaminolysis: We did not observe a significant increase of glutaminolysis in cancer vs normal. However, the metabolomic change of intermediate substrates suggests that the fueling role of glutaminolysis truly relieved the largely depleted carbon source in the TCA cycle.

Glutathione metabolism: Our analysis included three input sources of GSH, namely biosynthesis from glutamate in mitochondria, biosynthesis in cytosol and transport into mitochondria, and reduction from GSSG. We also considered two out-fluxes of GSH, namely biosynthesis of other amino acids and enzyme catalyzed oxidation of GSH to GSSG. We observed increased in-flux of GSH from glutamate and cytosolic biosynthesis, decreased out-fluxes of GSH, and accumulated GSH abundance in cancer vs normal. Noted, our flux estimation only covers enzyme catalyzed reaction of GSH to GSSG. Based on the observations, we speculate that majority of “accumulated” GSH predicted by scFEA are actually utilized for anti-oxidation, the flux of which was not included in the analysis.

Other branches: Our analysis also examined six branches of the central metabolism network. We observed an increase of nucleotide biosynthesis and transport of mitochondrial 2OG to cytosolic 2OG, decreased serine biosynthesis (except for KIRC and KIRP), fatty acid biosynthesis, and aspartate biosynthesis. We did not observe a significant flux from 2OG to immunosuppressive metabolite 2HG. A possible reason is that the reaction from 2OG to 2HG is catalyzed by mutated IDH1 or IDH2 enzymes, which were not covered by the current model.

The predicted fluxome and metabolomic changes are provided in **Supplementary Table S1**.

### Pan-cancer analysis suggests a convergent metabolic phenotype of human cancer

We conducted pan-cancer comparative of the predicted fluxome by using tSNE based dimensional reduction. **Figure 4A** show the 2D-tSNE plots of the analyzed samples from different cancer types derived by using the predicted flux distribution of the central metabolism network. **Supplementary Figure SX** list detailed distribution of each cancer and normal tissue types over the tSNE plot. We further conducted a K nearest neighbor clustering of the predicted flux based on their Euclidean distance and identified 12 clusters (**Figure 4B**). We have observed that (1) normal tissue types have distinct metabolic phenotype; (2) although majority of cancer tissues have different metabolic phenotypes compared to normal tissues, the fluxome of central metabolism in some cancer tissues such as breast and colon cancer are more similar to their matched normal tissues; (3) the fluxome of some cancer types are similar, such as breast, colon, kidney and lung cancer; and (4) head and neck cancer, kidney renal papillary cell carcinoma, prostate and thyroid cancer have distinct metabolic phenotypes.

**Figure 4.**
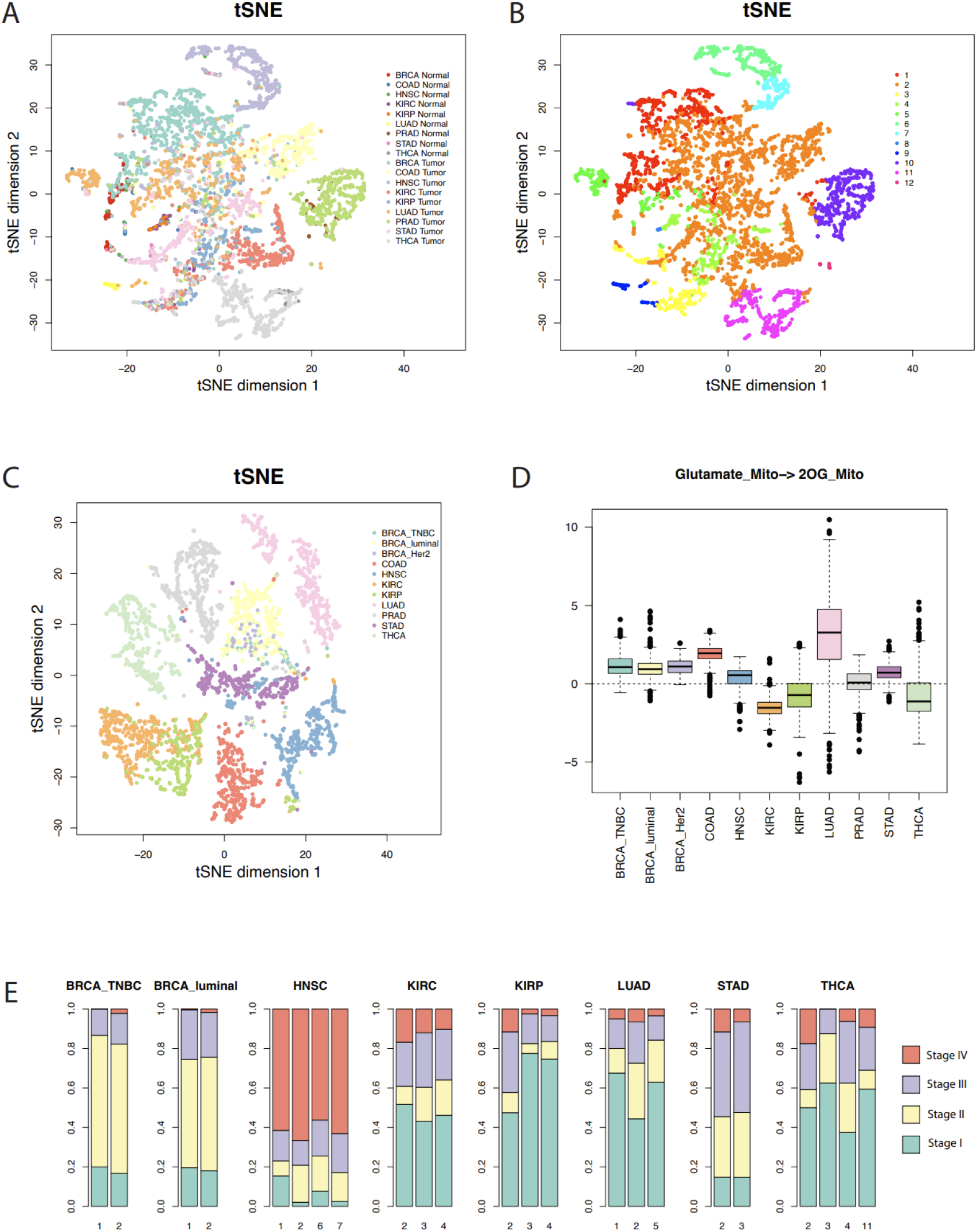
Metabolic phenotypes identified from pan-cancer analysis. (A-B) tSNE plots of the cancer and normal tissues derived by using predicted flux. (C) tSNE plot of the cancer tissues derived by using flux change between cancer vs adjacent normal controls. (D) Flux change between cancer vs adjacent normal controls of the reaction of mitochondrial glutamate to 2OG. (E) Distribution of cancer stages in each cluster with respect to cancer types.

Interestingly, we note a cluster (the cluster 2 in **Figure 4B**) consist large sets of breast, colon, kidney, lung, stomach cancer, normal breast and colon, and small sets of head and neck, prostate, and thyroid cancer samples, while the other clusters are either tissue or cancer type specific. **Supplementary Table S2** lists the distribution of cancer and normal tissue types in each cluster. We further conducted a second tSNE analysis of the fluxome in cancer normalized by their adjacent normal controls. Specifically, we computed the Z score of the flux of each module in a cancer sample against the flux of all normal controls of this cancer type and utilized the Z score profile for tSNE analysis (**Figure 4C**). Our results suggest that (1) normal tissues have distinct metabolic phenotypes, (2) cancer types have drastically different metabolic shifts, and (3) the different shifts happened to tissue specific metabolic phenotypes result in a converged metabolic phenotype through cancer types. **Figure 4D** illustrates the shifts of flux of mitochondrial glutamate to 2OG, a key step in glutaminolysis, which show a significant cancer or tissue type specificity. Similarly, majority of the reaction modules in the central metabolic network show different level of shifts through cancer types.

We further checked the distribution of cancer stage versus the identified clusters. The cluster 2, which was the converged metabolic phenotype through multiple cancer types, is consistently more enriched by the cancer of more advanced stages, for all the cancer types enriching to this cluster (**Figure 4E**). Hence, with the progression of cancer, metabolic shifts tend to converge metabolic phenotype to this cluster. Further analysis suggests that this cluster has increased glycolytic activity, lactate production, decreased TCA cycle, slightly increased glutaminolysis, and more saved glutathione for potential antioxidation. However, this cluster does not have the highest change in such metabolic shifts compared to other clusters. Hence, we speculate that the central metabolic system, biochemical condition, redox balance, and demand of energy and substrates of the cancer tissues in this cluster are more tuned.

Sun et al summarized 42 metabolic stress marker gene sets. We also examined the correlation between sample-wise gene expression level of the metabolic stress marker sets computed by single sample Gene Set Enrichment Analysis (ssGSEA) and predicted fluxome (**Figure 5**). As majority of the stress marker sets are biosynthesis related, the decreasing of which suggests elevated metabolic stress or unmet demand. We observed that the TCA cycle, biosynthesis of amino acids and fatty acids, and enzyme catalyzed reaction of GSH are positively correlated with the stress marker sets, while the glycolysis, lactate production, glutaminolysis, and antioxidation role of GSH have negative correlations. Our analysis identified biosynthesis favored (positive correlation) and un-favored (negative correlation) metabolic modules in the central metabolism network. Noted, glutaminolysis related modules, especially the import of glutamine and glutamate are unfavored in the cancer tissue with high biosynthesis, suggesting limited role of glutaminolysis in fueling biosynthesis of large molecules. An alternative explanation is that when redox reaction involved biosynthesis activity were suppressed, more glutamine and glutamate are needed by cancer cells to sustain sufficient amino acids.

**Figure 5.**
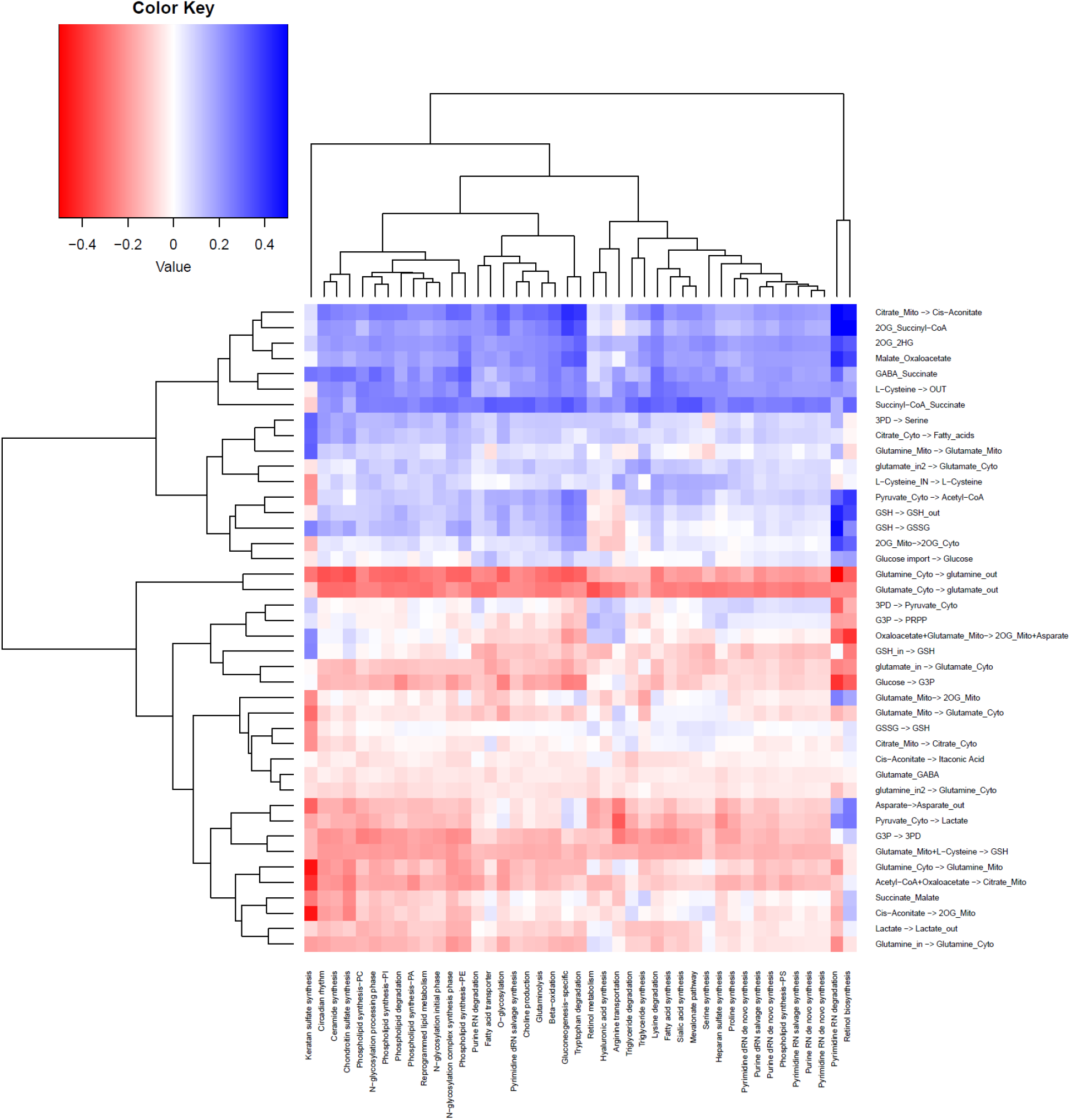
Correlation between the fluxome of central metabolic network and the expression level of 42 metabolic stress related modules.

## DISCUSSION

In this study, we reported the convergence of metabolic phenotypes in disease progression of multiple cancer types. This result is consistent with our previous observations on single cell RNA-seq data of the TME of melanoma and head and neck cancer, i.e., immune and stromal cells tend to have higher metabolic heterogeneity compared to cancer cells [2]. While Warburg effect has been identified as a common hallmark of almost all cancer types, heterogeneity in the means of energy production has rarely been studies, i.e., the choice between anaerobic glycolytic pathway and aerobic TCA cycle pathway. Our observations provide new theories of the trajectories of metabolic shift that occurred in oncogenesis process, that is, for a large set of solid cancer types, their metabolic phenotypes, determined by the flux distribution of the central metabolic pathways, tend to converge through the oncogenesis process. This indicates that it is the converged phenotype, rather than the path to convergence, that embodies the hallmark property of cancer progression. This “convergence” theory is rational as for cancer cells, the optimal flux distribution should allow cells to sustain fast cell proliferation rate and high fitness level under hostile and dynamic biochemical conditions in TME, which is independent of cancer types and tissue origins. This theory could also explain why metabolic inhibitors have not been very successful in cancer treatment. While different tumor tissues may converge to the same metabolic state, they may undergo different paths to such convergence. Hence, simply blocking a particular path may not stop the cancer progression to the desirable phenotype. A future direction is to comprehensively characterize the converged metabolic state that are most desirable for cancer progression, including the level of reaction rate and ratio of energy produced using different branches. Also, new computational measures are needed to identify the trajectory of metabolic shifts for each cancer and the distance of its current metabolic phenotype to the optimal metabolic state. Our current analysis suggests that genetic mutation is not enough to explain how heterogeneous paths are formed towards an optimal metabolic phenotype. We speculate the shifts and evolution of metabolism is triggered by the co-evolution between cancer and the intra- and inter-cellular biochemical condition within its TME and facilitated by epigenetic regulations.

Numerous computational analyses have been proposed to study metabolic variations in cancer and other systems [8–13]. However, while substantial efforts have been paid on reconstructing metabolic pathways, a fundamental question remaining un-addressed is how metabolic activities differ among cells of different morphological types, physiological states, tissues, or disease backgrounds that have the same genetic constitutions. Although transcriptomics or metabolomics experiments have been utilized to characterize metabolic alterations in diseases [14, 15], existing analysis tend to portray the average change of intermixed and heterogeneous cell subpopulations within a given tissue [16–18]. This makes it impossible to further study the metabolic heterogeneity and cell-wise flux changes in a complex tissue, in which cells are well understood to rewire their metabolism and energy production in response to varied biochemical conditions [19–22]. Compared to other well studied biological mechanisms, such as immuno-response or transcriptional regulatory activity, there is substantial gaps in characterizing metabolic changes using omics data, with tailored systems biology models and statistical metrics.

To the best of our knowledge, our recently developed scFEA is the first and only method to estimate cell-wise metabolic flux and metabolomic changes by approximating the underlying dynamical systems models [23]. Our analysis demonstrated the feasibility to decipher the cause and impact of metabolic variations by using multi-omics data in an explicit way, namely data-driven and AI-empowered systems biology. On the other hand, in our preliminary study, we identified three major remaining challenges_in leveraging the high-resolution multi-omics data and the stochiometric relations of the metabolic network that need to be solved to best characterize the dynamics and context specificity of metabolic activities: (1) *How to reconstruct disease, tissue, and cell group specific metabolic network?* A complex tissue microenvironment may be constituted by cells of different metabolic abnormalities, heterogeneous metabolic networks, varied preferences, and dependencies [24–28]. Mapping multi-omics data to a common and static metabolic network precludes the discoveries of hidden and dynamic relationships among the metabolic units, making it impossible to identify the key players in disease tissue or cells, and to predict vulnerability of a particular phenotype to a certain metabolic factor. (2) *How to accurately estimate metabolic flux and identify the key causes of metabolic variations by using multi-omics data?* A big gap in metabolic modeling is how to map diverse data types onto quantitative metabolic models in order to elucidate more thoroughly the metabolic fluxome, and hence to achieve functional characterization and accurate quantification of all levels of metabolic activities and their interactions [29–31]. Although our recent progress and other studies provide a preliminary solution, no existing method can effectively handle the heterogeneity of directions of highly reversible reactions and imbalance of intermediate metabolites among cells within a disease microenvironment. (3) *How to comprehensively define and assess the mechanisms and representation forms of metabolic variations on multi-omics data?* Metabolic variations happen on different levels, such as gene, enzyme, metabolite, network structure or flux (kinetic models). It remains unsolved on how to design valid metrics and statistical models to quantify the true impact of such variations on the context specific metabolic activity.

Noted, our study and method demonstrated a prototype of a new research direction, namely “data-driven and AI empowered systems biology”. For given omics data and a to-be-studied biological process, the underlying goal is to identify a mathematical model that could not only quantify the biological process but also approximate its dynamic property over the data. The established model should leverage the coherency to the physical or chemical laws of the system and the goodness of fitting to the data. Compared to conventional differential equation-based systems biology model, our approach does not rely on pre-assessed kinetic parameter, hence is not limited by the reductionist paradigm can be applied to characterize a relatively large system, such as the central metabolism network, in complex disease.

## METHODS

### Data used in this study

<u>TCGA transcriptomic data</u>: TCGA RNA-seq v2 FPKM data of the nine cancer types (11 subtypes) were retrieved from the Genomic Data Commons (GDC) data portal using TCGAbiolinks [32]. Table 4 lists the names of the cancer types along with the information of the numbers of cancer and control samples. FPKM values were converted to Transcripts per Million (TPM) values as the latter is more stable across samples. Clinical data were obtained in XML format from GDC and parsed with an in-house script. GENCODE gene annotations used by the GDC data processing pipeline was downloaded directly from the GDC reference files webpage.

### Main methods

<u>scFEA</u>: We have directly applied our scFEA method on the TCGA and the two scRNA-seq data against the iron metabolic map. While the details of the method are given in [33], we outline the key ideas of the algorithm. The inputs to scFEA are a gene expression data and a factor graph-based representation of the metabolic map. Let *FG*(*C*^1×*K*^, *RM*^1×*M*^, *E* = {*E*_*C*→*R*_, *E*_*R*→*C*_}) be a given factor graph, where *C*^1×*K*^ = {*C_k_,k* = 1, …,*K*} is the set of *K* metabolites, *RM*^1×*M*^ = {*R_m_,m* = 1, …,*M*} the set of *M* metabolic reactions (represented as a rectangle in Figure 1A), *E*_*C*→*R*_ and *E*_*R*→*C*_ represent direct edges from reaction *R_m_* to metabolite *C_k_* and from metabolite *C_k_* to reaction *R_m_*, respectively. For the *k* -th metabolite *C_k_*, define the set of reactions consuming and producing *C_k_* as 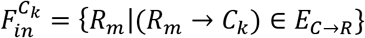 and 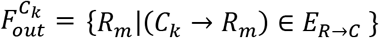, which is derived from the stoichiometric matrix of the given metabolic map. For an RNA-seq dataset with *N* cells, denote *Flux_m,j_* as the flux of the *mth* reaction in the cell *j,j* = 1 …*N*, and *F_j_* = {*Flux*_1,*j*_,…, *Flux_M,j_*} as the whole set of the reaction fluxes. Denote 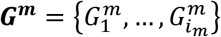 as the genes associated with the reactions in *R_m_*, and 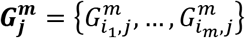 as their expressions in sample *j*, where *i_m_* is for the number of genes in *R_m_*.

We model 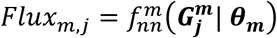 as a multi-layer fully connected neural network with the input 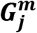, where ***θ_m_*** represents the parameters of the neural network. Then the ***θ_m_*** and cell-wise flux *Flux_m,j_* can be solved by minimizing the following loss function L, where *λ* serves as a hyperparameter:

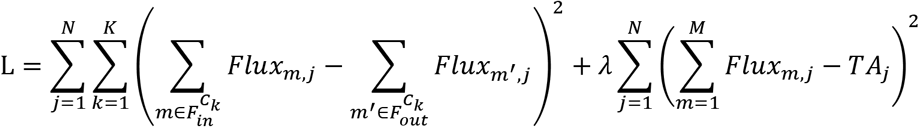

where *TA_j_* is a surrogate for the total metabolic level of cell *j*, which is assigned to a constant or total expression of all the metabolic genes in *j*.

It is noteworthy that this formulation defines a new graph neural network architecture for flux estimation over a factor graph, where each variable is defined as a neural network of biological attributes, i.e., the genes involved in each reaction; and information aggregation between adjacent variables is constrained by the imbalance between the in- and out-lux of each metabolite.

<u>tSNE analysis</u> was conducted by using Rtsne v0.16 R package against full fluxome profile with default parameters.

<u>K-near neighbor clustering</u> was conducted by using Seurat v3 R package against top 12 principal components. The number of clusters is determined by default settings in the Seurat package.

ssGSEA: We applied ssGSEA2.0 R package to estimate the levels of the selected RMs on individual samples [34, 35]. The ES score computed by ssGSEA was utilized to represent the level of each RM. Gene sets of the RMs were collected and annotated in our previous work [33].

<u>Statistical test of differential analysis</u>: We have utilized Mann Whitteny test for all differential analysis, including differential gene expression analysis and the difference of predicted flux.

## ACKNOWLEDGEMENTS AND FUNDING RESOURCES

The work was supported by NSF DBI IIBR 2047631, NSF IIS 2145314, American Cancer Society RSG-22-062-01-MM, and NCI 5P30CA082709-22. C.S. and Z.C. want to thank the support from Indianapolis SEED and STEM project and project director Mr. Elmer Sanders.

